# Keep your hands apart: independent representations of ipsilateral and contralateral forelimbs in primary motor cortex

**DOI:** 10.1101/587378

**Authors:** Ethan A Heming, Kevin P Cross, Tomohiko Takei, Douglas J Cook, Stephen H Scott

## Abstract

It is generally accepted that each cortical hemisphere primarily drives the opposite side of the body. Yet, primary motor cortical (M1) activity has been robustly correlated with both contralateral and ipsilateral arm movements. It has remained unanswered as to why ipsilaterally-related activity does not cause contralateral motor activity. Here we apply multi-joint elbow and shoulder loads to the left or right arms of monkeys during a postural perturbation task. We show that many M1 neurons respond to mechanical disturbances applied to either the contra- or ipsilateral arms. More neurons respond to loads applied to the contralateral arm with response magnitudes that were ~2x as large and had onset times that were ~10ms earlier. However, in some cases, neurons exhibited large and earlier responses to loads applied to the ipsilateral arm than loads applied to the contralateral arm. Similar effects were observed when the monkeys were maintaining postural control well after the load had been applied. Importantly, we show that the load preference to one arm has little predictive power on a neuron’s preference in the opposite arm. Furthermore, we found contralateral and ipsilateral neural activity resided in orthogonal subspaces allowing for a weighted sum of neural responses to extract the contralateral activity without interference from the ipsilateral activity, and vice versa. These data show how activity in M1 unrelated to downstream motor targets can be segregated from downstream motor output.

## INTRODUCTION

Several lines of research demonstrate that primary motor cortex (M1) is principally involved in controlling movements of the contralateral side of the body. Anatomically, greater than 90% of corticospinal projections target the contralateral side (Dum and Strick, 1996; Brösamle and Schwab, 1997; Lacroix et al., 2004; Rosenzweig et al., 2009). Most of the 10% that project ipsilaterally bifurcate and synapse bilaterally, with few being thought to synapse onto purely ipsilaterally targets (Rosenzweig et al., 2009). While there are many direct projections from M1 to contralateral limb muscles (Bennett and Lemon, 1996, 1996; McKiernan et al., 1998; Smith and Fetz, 2009), there are no monosynaptic projections from M1 to ipsilateral muscles (Soteropoulos et al., 2011). Stimulation in M1 largely produces contralateral movements and occasionally bilateral movements (Montgomery et al., 2013). These studies highlight that most M1 descending projections principally target the contralateral side of the body.

Early studies that recorded M1 activity during ipsilateral movement suggested M1 was largely insensitive to ipsilateral movement (Tanji et al., 1988). However, several follow-up studies highlight substantial M1 activity during ipsilateral motor behaviours (Kermadi et al., 1998; Cisek et al., 2003). Donchin et al., (2002) examined macaque M1 neuronal activity in relation to ipsilateral and contralateral reaching movements. They found that only 34% of cells responded to only contralateral movements, whereas 19% responded to only ipsilateral hand movements, and 46% responded to both. Functional magnetic resonance imaging studies have also found considerable M1 activity related to ipsilateral movement (Cramer et al., 1999; Gallivan et al., 2011). Kobayashi et al. (2003) recorded BOLD responses while subjects completed rhythmic index finger tapping and found activity in the ipsilateral sensorimotor cortex. Diedrichsen et al. (2013) found the cortical area associated with the ipsilateral digit overlapped with its representation for the corresponding contralateral digit.

An obvious question is why does ipsilateral activity in M1 not lead to contralateral limb movement? Recently, it has been shown that when planning for a reach, motor cortex generates preparatory activity that resides in an orthogonal subspace to the activity during movement (Churchland et al., 2012; Kaufman et al., 2014; Elsayed et al., 2016). This can occur if the net change in population activity cancels in the movement dimensions (Druckmann and Chklovskii, 2012). For example, consider the toy example of 2 neurons that synapse with equal, excitatory synapses onto an alpha-motoneuron. If both neurons increase their firing rate, the result will lead to excitation in the alpha-motoneuron and thus movement (movement subspace). Conversely, if one neuron increases and the other neuron decreases its firing rate equally, the net effect on the alpha-motoneuron will be zero leading to no movement (null subspace). Examining the movement and null patterns in state space, where each axis represents the firing rate of a neuron, identifies that these two patterns are orthogonal to each other. This strategy can allow motor cortex to perform computations necessary for the upcoming movement without causing movement. This was demonstrated by Stavisky et al., (2017) where early visual feedback about a displacement to the cursor position was isolated from the subsequent corrective motor response. Also, this strategy can be used to engage different spinal cord pathways that lead to a motor output (Miri et al., 2017). We hypothesize a similar process may exist for M1 activity related to the ipsilateral limb.

We used a postural perturbation task to explore M1 activity related to the contralateral and ipsilateral limbs. We found ~55% of neurons were active when loads were applied to the contralateral and ipsilateral limbs. However, contralateral loads tended to evoke neural responses that were twice as large as responses for ipsilateral loads. Furthermore, contralateral loads evoked changes in neural activity ~10ms earlier than ipsilateral loads. Lastly, we found contralateral and ipsilateral activity resided largely in orthogonal subspaces suggesting a mechanism of how motor cortex can sequester the ipsilateral activity without causing movement.

## METHODS

### Animal and apparatus

Studies were approved by the Queen’s University Research Ethics Board and Animal Care Committee. Two non-human primates (NHP, *macaca mulatta*) were trained to perform a postural perturbation task similar to those used in our previous studies (Herter et al., 2009; Omrani et al., 2014; Heming et al., 2016) using a KINARM exoskeleton robot (BKIN Technologies, Kingston, Canada; (Scott, 1999)). On each trial, the monkey maintained its right or left hand, represented by a white cursor (0.5cm diameter), at a stationary virtual target (0.8cm diameter, red for right, blue for left, luminance matched). These targets were positioned at locations approximately in front of each shoulder, with the shoulder at 30° forward flexion and the elbow at 90° flexion (Fig. 1A). Only one target was shown at a time, thus only one arm was used in a trial. After an initial unloaded hold period of 500-1000ms, a random flexion or extension step load was applied to the shoulder and/or elbow. After the load was applied, the monkey had 1000ms to return their hand to the target. Once inside the target, the monkey had to maintain its hand inside the target for 1000-1500ms to receive a water reward.

**Figure 1:**
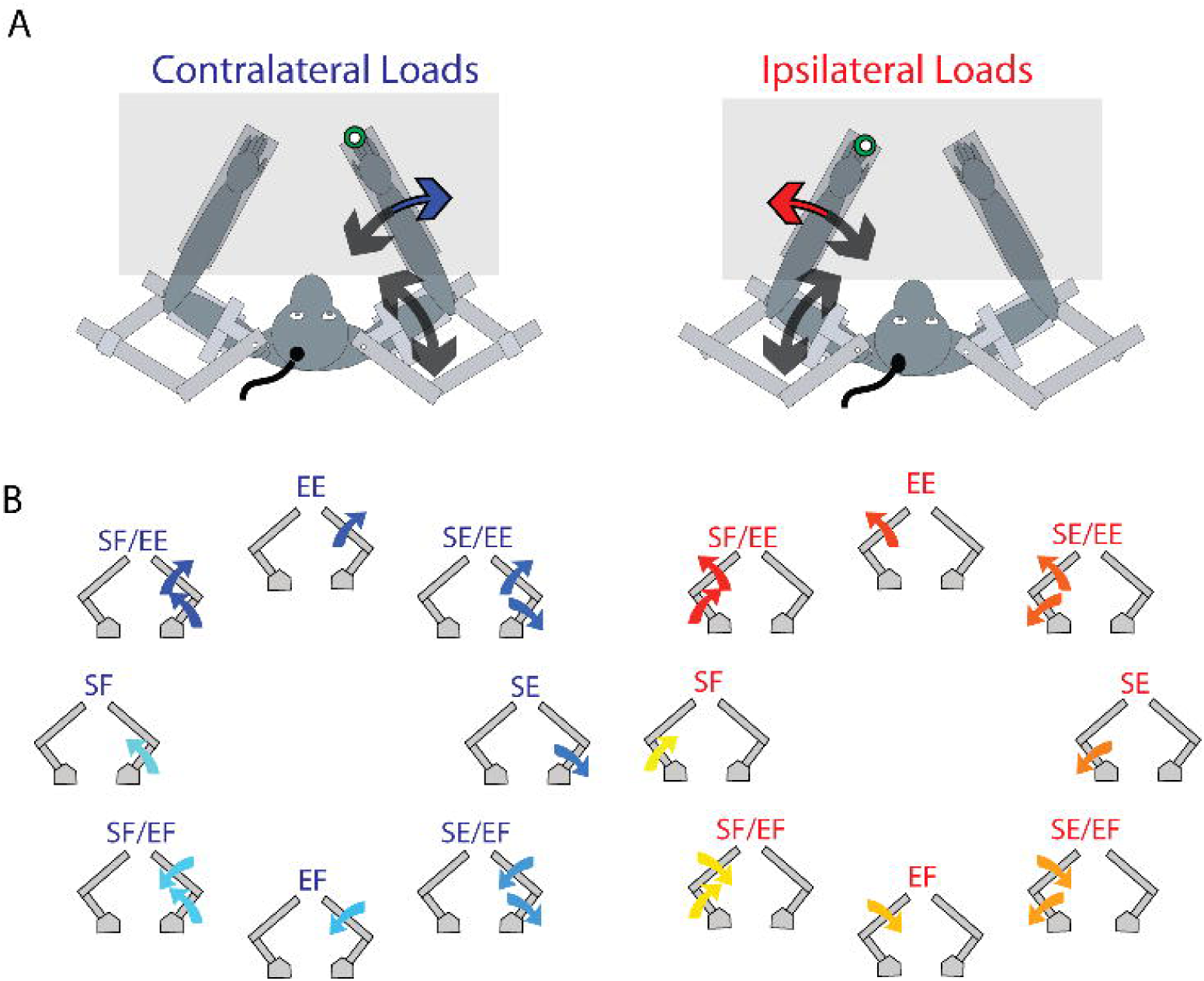
Experiment Set-Up. A) The monkey’s left and right arms were supported by the exoskeleton. Monkeys were trained to return their hand to the goal target (green) when mechanical loads were applied to the right (contralateral blue arrow) and left (ipsilateral red arrow) hand. Visual feedback of their hand was presented as a white cursor. B) For contralateral and ipsilateral loads, combinations of flexion and extensions torques were applied to the shoulder and elbow joints (arrows).

Eight load conditions were used for each arm, consisting of torques that caused elbow flexion (EF), elbow extension (EE), shoulder flexion (SF) or shoulder extension (SE), or the four multi-joint combinations of torques (SF/EF, SF/EE, SE/EF, SE/EE, Fig. 1B). The magnitude of the torques were 0.20 Nm for single joint loads, and 0.14 Nm at each joint for multi-joint torques. All torque conditions were completed in random order, comprising one block. A minimum of 10 blocks were completed.

### Neural, EMG and kinematic recordings

After training, monkeys underwent surgery to implant 96-channel Utah Arrays (Blackrock Microsystems, Salt Lake City, UT) into the arm region of M1. Surgeries were performed under aseptic conditions and a head fixation post was also attached to the skull using dental cement. Spike waveforms were sampled at 30kHz and acquired using a 128-Channel Neural Signal Processor (Blackrock Microsystems, Salt Lake City, UT). Spikes were manually sorted offline (Offline Sorter, Plexon Inc., Dallas TX) using a space spanned by the top two principal components and the peak-trough voltage difference. Only well isolated single-units were used for analysis. For Monkey P we also implanted a chamber above the right arm area of M1 and neurons were recorded via single electrode over multiple recording sessions using standard techniques (Herter et al., 2009).

We recorded intramuscular electromyographic (EMG) activity from Monkey P by inserting two thin wires into the muscle belly of brachioradialis, lateral and long head of triceps, long head of the biceps and pectoralis major. Signals were digitized at 1KHz.

Joint angles, velocities, and accelerations for both arms, were recorded at 1 kHz by the Neural Signal Processor. All offline analysis was performed using custom MATLAB scripts (The MathWorks, Inc., Natick, Massachusetts, United States).

### Kinematic analysis

Kinematics were filtered using a 3^rd^ order high-pass Butterworth filter with a cutoff frequency of 10Hz. Integrated hand speed was calculated by summing the hand speed from 0 to 300ms after the perturbation. A paired t-test was used to assess if the magnitude of the integrated hand speed was larger when contralateral or ipsilateral loads were applied.

### EMG analysis

EMG signals were aligned to the perturbation onset and filtered using a 6^th^ order bandpass Butterworth filter with a frequency range of 20-200Hz. Signals were then rectified and further smoothed with a 6^th^ order low-pass Butterworth filter with a cutoff frequency of 100Hz. We tested for load sensitivity by applying a two-way ANOVA with time epoch (levels: baseline and perturbation) and load combination (levels: 8 load combinations) as factors. Samples were deemed significant if a main effect of time or an interaction effect was significant (p<0.05). Group average signals were generated by finding the muscle’s preferred load direction and normalizing the EMG signal by the mean activity in the perturbation epoch.

### Spike trains smoothing and epochs

Spikes were convolved with an asymmetric kernel approximating a post-spike potential (1ms rise time and 20ms fall time) (Thompson et al., 1996). This kernel is causal, meaning it only influences the firing rate after rather than before the spike, which provides a better estimate of timing onsets than non-causal Gaussian kernels. We constructed two trial-averaged histograms, one that was aligned to the perturbation onset and spanned the first 300ms after the perturbation (perturbation epoch) and one that spanned the last 1000ms of the trial (steady-state epoch).

### ANOVA analysis

For each neuron, we binned its firing rate into a baseline (200ms before load onset till load onset) and perturbation (first 300ms after load onset) epoch for each context (i.e. contralateral and ipsilateral) and load combination. A three-way ANOVA was then applied with timing epoch, context and load combination as factors. We classified neurons as load sensitive if they had a significant main effect for time, or any interaction effects with time (p<0.05).

### Preferred load direction and timing onset

For each neuron we time averaged the firing rates within each epoch and subtracted off the mean signal. We then fit a planar model (MATLAB *regress*) that mapped a neuron’s firing rate to the applied loads. Rayleigh unimodal and bimodal r-statistics were calculated for the population distribution of preferred load directions (Batschelet, 1981; Lillicrap and Scott, 2013). We compared our results to a bootstrapped distribution generated by randomly sampling angles from a uniform distribution spanning 0-360°. We matched the number of angles we sampled with the number of neurons in our population of interest. We then calculated the unimodal and bimodal r-statistics for the resulting bootstrap distribution and repeated this procedure 1000 times. Significance was assessed by calculating the percentage of times we found the bootstrapped sample had a larger r-statistic than our neuron population.

We estimated the timing onsets by first finding the load combination that generated the absolute largest change in firing rate from baseline (binned activity from 200ms before load was applied till load onset) during the perturbation epoch. We then calculated the trial average by including trials from the 2 spatially adjacent load combinations to improve onset estimate (Herter et al., 2009). Onsets were calculated by finding the first time point that exceeded baseline activity by 3 standard deviations and remained above the threshold for 20 consecutive milliseconds.

### Correlation matrices and PCA

We used previously established methods for our correlational and PCA analyses (Elsayed et al., 2016; Miri et al., 2017). First, we down sampled the firing rate of each neuron by sampling every 10ms. We then soft-normalized the contralateral and ipsilateral activity of each neuron by its maximum firing rate plus 5 spikes/s. For each context (contralateral vs ipsilateral loads), we subtracted off the mean neural activity across the eight perturbation types for each time bin. For contralateral and ipsilateral activity, we constructed matrices C and I ∈ *R^NxCT^* where N is the number of neurons, C the number of mechanical loads (8) and T the number of time points (perturbation epoch: 30 time points after down sampling).

Correlation matrices were generated by comparing each neuron’s correlation with all other neurons within a context (ie. contralateral or ipsilateral activity). We revealed structure underlying the correlation matrices by first choosing a threshold of 0.4 and assigning 0 or 1 for each element in the correlation matrix that was above or below this threshold. We then used the MATLAB function *symamd* to find a column and row permutation that clustered nonzero elements. These permutations were then applied to the contralateral and ipsilateral correlation matrices.

In order to observe how correlations between neurons change between the two contexts, we computed the correlation between each possible pair of neurons during each context. This yielded two correlation coefficients for each pair of neurons, one for each context. We then calculated the absolute change between the two correlation coefficients for each neuron pair. We compared the resulting distribution across all neuron pairs with a bootstrapped null distribution (Figure 7D and K), as previously described (Miri et al., 2017). This distribution is generated from the assumption that the absolute change in correlation between the two contexts are due to averaging over a finite number of trials. For each neuron, we randomly assigned each of its trials during one of the contexts to two separate groups and generated trial-averaged firing rates. Within each group we calculated all pairwise correlation coefficients between neurons and then calculated the absolute change in coefficients between groups. This was repeated 1000.

PCA was performed on matrices C and I using singular value decomposition. We selected the top 10 principal components for each context and calculated the amount of contralateral and ipsilateral variance each projection could account for (VAF).

We determined how aligned the top-ten contralateral and ipsilateral principle components were by the alignment index, as described previously (Elsayed et al., 2016):

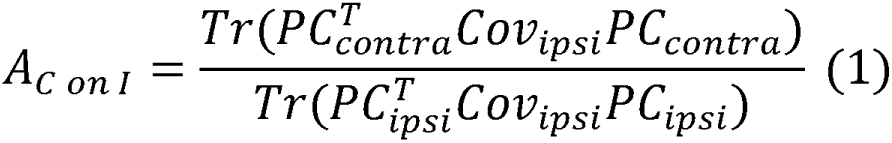

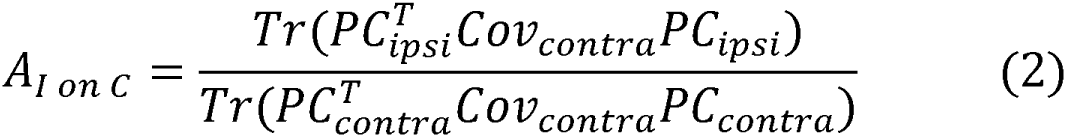

where *PC_contra_* and *PC_ipsi_* ∈ *R*^10*xCT*^ are matrices containing the top-ten contralateral and ipsilateral principal components, *Cov_contra_* and *Cov_ipsi_* are the contralateral and ipsilateral covariance matrices (*R^nxn^*), and Tr is the trace operator. Equation 1 reflects a ratio of how much variance the top-ten contralateral principal components could explain of the ipsilateral activity with how much the top-ten ipsilateral principal components could explain of the ipsilateral activity (the max variance any ten linear projections could capture). This metric was also computed using the contralateral activity (Equation 2). The alignment index takes a range from 0, which indicates the contralateral and ipsilateral principal components are perfectly orthogonal, to 1 which indicates the contralateral and ipsilateral principal components are perfectly aligned.

We assessed if the alignment indices were significant by a bootstrap procedure as described in Elsayed et al., (2016). Initially, PCA was performed on the data matrix generated by concatenating the contralateral and ipsilateral matrices, C and I. Two sets of ten components were randomly sampled from the data matrix with a probability of selection weighted by the amount of variance for that component. This procedure was repeated 1000 times to generate a null distribution.

### Orthogonalization

We found an orthogonal set of projections that captured a significant amount of contralateral and ipsilateral activity by using a joint optimization technique as described in Elsayed et al., (2016). This technique seeks a set of projections that maximizes the amount of variance explained for contralateral and ipsilateral activity while constraining the projections to be orthogonal. The cost function for this optimization was

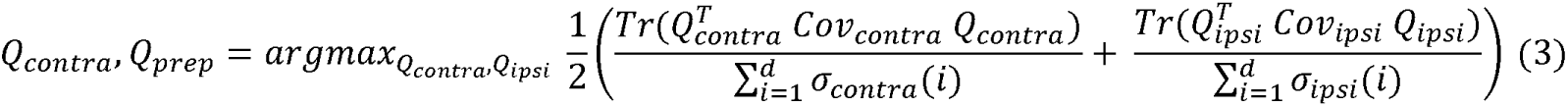

where d is the number of dimensions, *σ_contra_*(*i*) and *σ_ipsi_* (*i*) are the i-th singular values of the contralateral and ipsilateral covariance matrices, and *Q_contra_*, *Q_ipsi_* are matrices ∈ *R^Nxd^* composed of the orthonormal components. With the orthonormal constraints on *Q_contra_*, *Q_ipsi_* the problem reduces to an optimization problem on a Steifel manifold which we found using Manopt for MATLAB (Boumal et al., 2014; Cunningham and Ghahramani, 2015). The resulting projections *Q_contra_*, *Q_ipsi_* were then used to reduce the contralateral and ipsilateral activity matrices C and I. For presentation purposes, we first applied PCA before plotting to order the projections from largest to smallest VAF, and we also smoothed the time series with a 3^rd^ order low-pass Butterworth filter with a cut-off frequency of 25Hz. However, all analyses utilized the original, unsmoothed time series.

The relative difference between the contralateral and ipsilateral time series in the orthogonal contralateral dimensions was calculated as

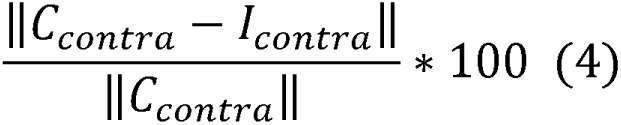

where *C_contra_* and *I_contra_* are the projections on the contralateral dimensions for the contralateral and ipsilateral activity, respectively. In order to estimate the variability with this metric, we bootstrapped ipsilateral trials and calculated the relative difference 1000 times. A similar calculation was computed for the ipsilateral projections.

## RESULTS

### Kinematics and EMG

Monkeys were trained to keep their right or left hand in a target and return their hand to the target after mechanical loads were randomly applied to the shoulder and/or elbow joints. Figure 2A-B shows Monkey P’s hand motion following mechanical loads that were applied to the contralateral arm only (right arm). The load caused the hand to deviate ~1cm from the starting position before the monkey returned the hand to the target. In the ipsilateral arm (left arm) we observed very little hand motion when the loads were applied to the contralateral arm. When loads were applied to the ipsilateral arm only, the ipsilateral arm deviated ~3cm from the starting position while the contralateral arm showed little hand motion (Figure 2C and D).

**Figure 2:**
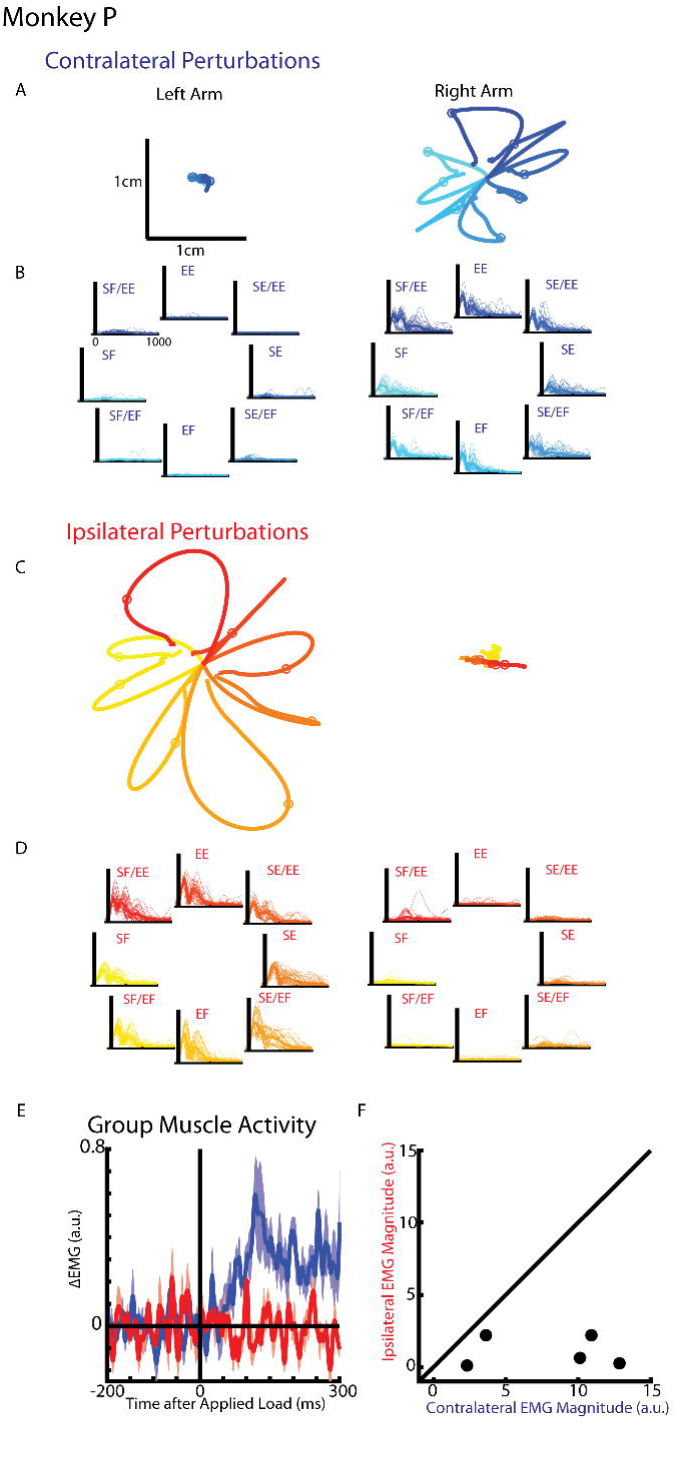
Hand kinematics when loads applied to the contralateral or ipsilateral arms. A) Average hand paths for the left and right hand from Monkey P when loads were applied to the contralateral arm (right arm). Circles indicate hand position 300ms after the load was applied. B) Single trial (thin lines) and average (thick) hand velocities when contralateral loads were applied. Black vertical line marks the onset of the load. C-D) Same as A-B for ipsilateral loads.

We compared the integrated hand speed of the contralateral and ipsilateral limb when the loads were applied. For the contralateral limb, the ipsilateral load caused motion that was a fraction of the motion caused by the contralateral load (mean for Monkey P: 6.6%, Monkey M: 10.3%) and was significantly smaller (paired t-test: Monkey P: t(7)=9.5, p<0.001, Monkey M: t(7)=9.9 p<0.001). Likewise, a similar reversal was found for the ipsilateral limb with the contralateral load causing motion that was only a fraction of the hand motion caused by the ipsilateral load (Monkey P: 9.9% t(7)=9.9 p<0.001, Monkey M: 12% t(7)=15.6 p<0.001). Similar results were found when we quantified the maximum hand speed as well as when we increased the interval of the time epoch (data not shown).

For Monkey P we recorded intramuscular muscle activity from shoulder and elbow extensors. Figure 2E shows the group average muscle response when contralateral and ipsilateral loads were applied. A clear increase in muscle activity can be observed in <50ms when contralateral loads were applied, however there was little change in muscle activity when ipsilateral loads were applied. Figure 2F compares the change in muscle activity from baseline in the perturbation epoch for the contralateral and ipsilateral loads. Most samples lie near the contralateral axis indicative of a larger contralateral than ipsilateral response. We applied a two-way ANOVA with time (levels: baseline, perturbation) and load direction (levels: 8 load combinations) as factors to the contralateral and ipsilateral muscle activity. All muscle samples had a significant main and/or interaction effect to the contralateral load whereas none were significant for the ipsilateral loads.

### Neural Recordings

From Monkey P and M, we recorded 92 and 130 neurons from M1, respectively. We first determined if each neuron was sensitive to the loads by applying a three-way RM ANOVA with time epoch (levels: baseline and perturbation epoch), context (levels: contralateral and ipsilateral) and load direction (8 levels: each load combination) as factors. We found 90 and 91 neurons in Monkey P and M had a significant main or interaction effect with time indicating a load sensitivity. From the load sensitive neurons, we then determined each neuron’s preferred load direction by regressing a plane of each neuron’s average firing rate (0-300ms epoch after the applied loads) with each load combination. Figure 3 shows average firing rates of 4 exemplary neurons for both contralateral and ipsilateral loads. Figure 3A shows the firing rate of an exemplar neuron that had a significant planar fit for the contralateral and ipsilateral loads. The neuron had a preferred load direction that was oriented towards elbow extension and shoulder flexion for contralateral loads and elbow flexion and shoulder extension for ipsilateral loads. Figure 3B shows a second exemplar neuron with preferred load directions in the opposite direction as the previous neuron for the contralateral and ipsilateral loads. Figure 3C and D show two neurons that had a significant fit to their firing rates for contralateral only (C) and ipsilateral only (D) loads, respectively.

**Figure 3:**
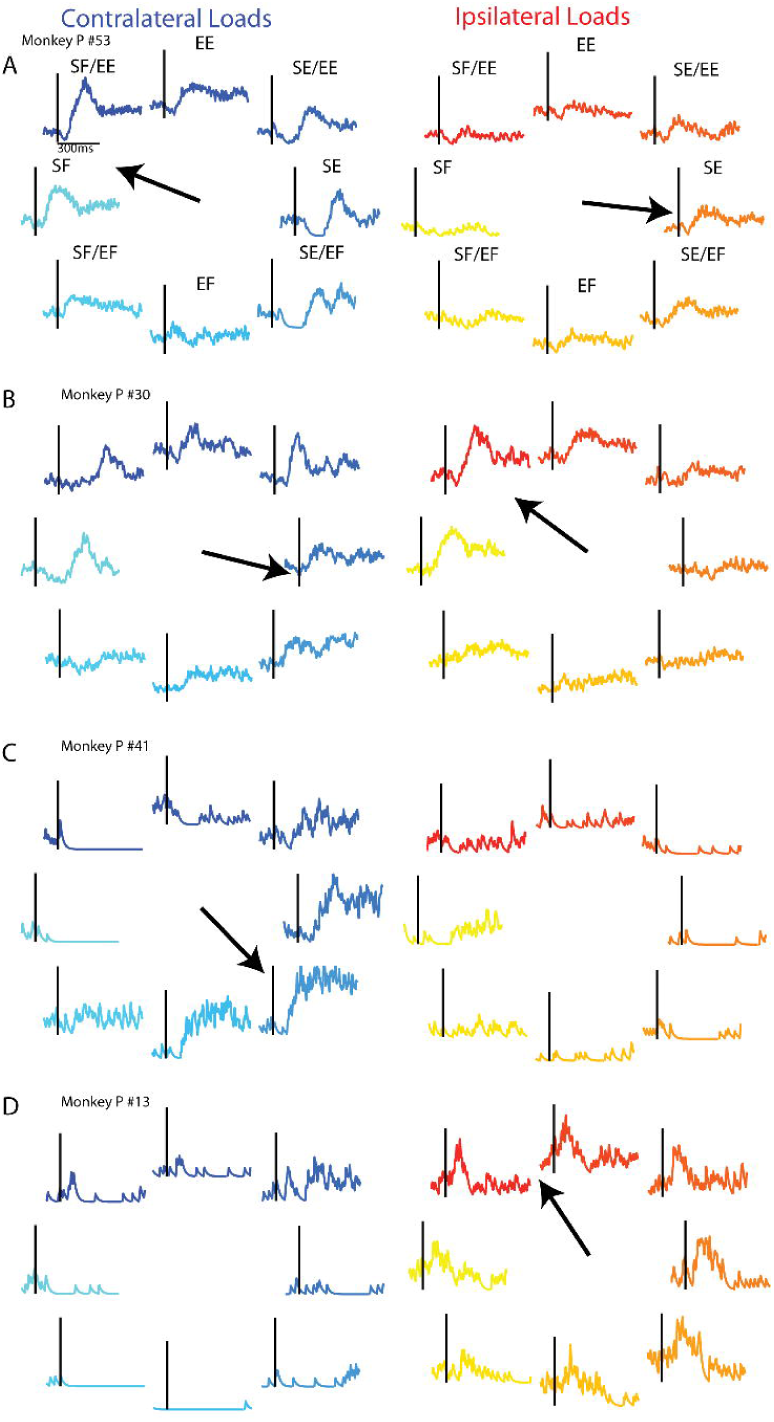
Exemplary neuron responses when contralateral and ipsilateral loads were applied. A) A neuron that had a significant plane for the contralateral (right) and ipsilateral loads (left). Arrows denote neuron’s preferred load direction. Vertical line in each panel denotes the time the load was applied to the arm. Eight load combinations were applied to each arm and are displayed following format in Figure 1. B) A second neuron that had a significant fit for the contralateral and ipsilateral loads. C) A neuron which had a significant plane fit for contralateral loads only. D) A neuron which had a significant plane fit for ipsilateral loads only.

For Monkey P/M we found 48/60% of load sensitive neurons also had significant planar fits for contralateral and ipsilateral loads, whereas 39/18% neurons had a significant fit for contralateral loads only and 11/11% neurons had a significant fit for ipsilateral loads only.

When we examined the steady-state (last 1000ms of trial), we found 51/53% neurons had significant planar fits for both loads for Monkey P/M, whereas 30/26% neurons had a significant planar fit for contralateral loads only and 16/13% neurons had a significant planar fit for ipsilateral loads only.

Figure 4A,B,F,G shows the distribution of the preferred load direction of the population of neurons for the contralateral and ipsilateral loads during the perturbation epoch (0-300ms) and the steady-state (last 1000ms of the hold period). For Monkey P and M, we found significant bimodal distributions for both contexts and time epochs when all load sensitive neurons were included. The major axis of the bimodal distributions was in the quadrants for shoulder extension/elbow flexion and shoulder flexion/elbow extensions (Figure 4A and B) quadrants 2 and 4, blue line), consistent with our previous results (Cabel et al., 2001; Herter et al., 2009). Analysis of the difference between the neuron’s tuning during the perturbation and steady-state epoch revealed a significant unimodal distribution with a major axis near zero (Figure 4D, E, I, J).

**Figure 4:**
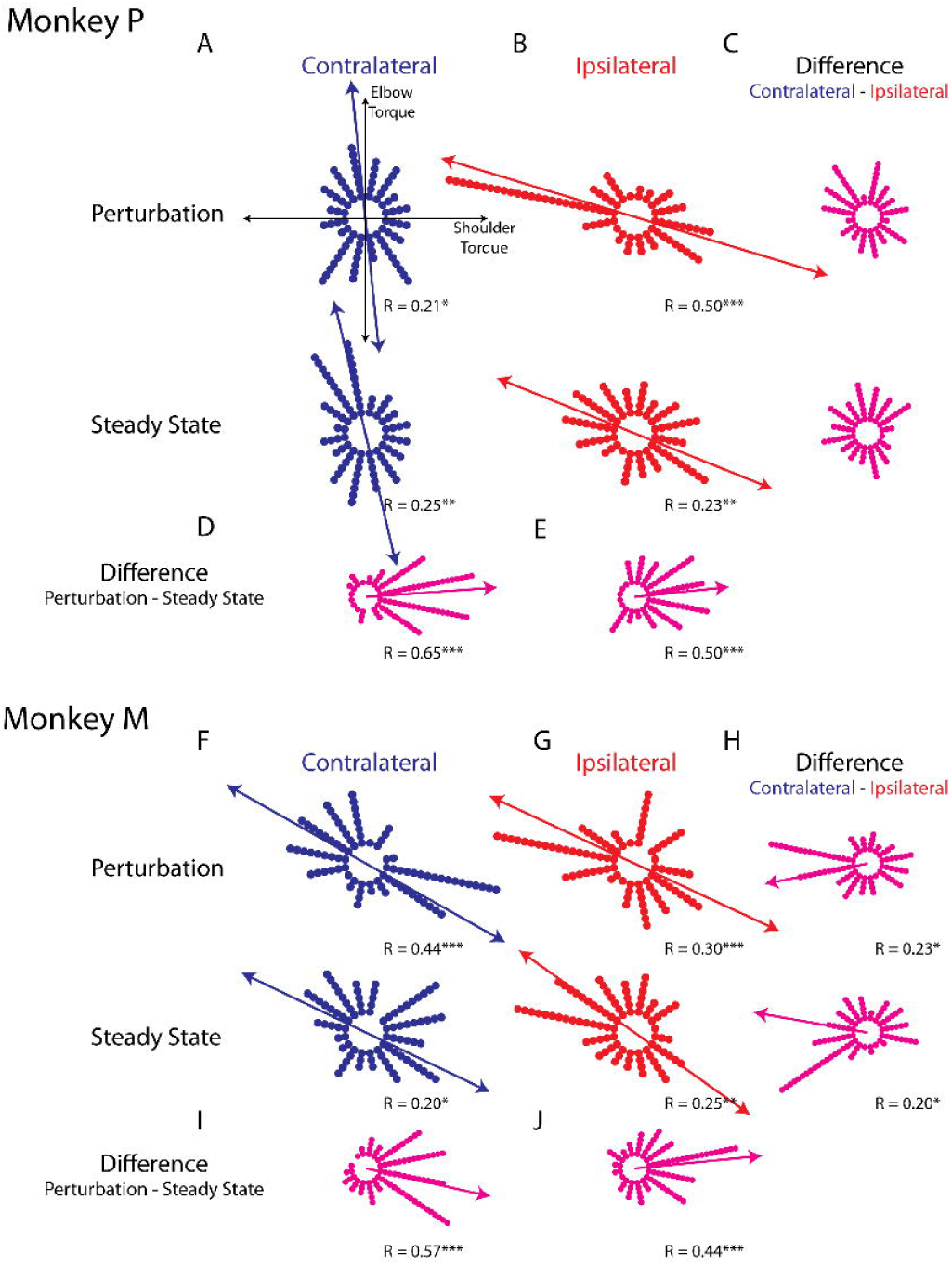
Tuning distributions in joint torque space. A) Polar histograms showing the distribution of tuning curves plotted in joint torque space for the perturbation (top) and steady-state (bottom) epoch when contralateral loads were applied. R reports the Rayleigh statistic. Blue arrow shows the major axis of the bimodal distribution. B) Same as A for the ipsilateral load. C) Polar histogram showing the change in tuning between the contralateral and ipsilateral loads. D) Polar histogram showing the change in tuning between the perturbation and steady-state epochs for contralateral loads. Magenta arrows indicate major axis for the unimodal distribution. E) Same as D for ipsilateral loads. F-J) Same as A-E for Monkey M.

When we examined the difference between each neuron’s tuning between the contralateral and ipsilateral loads, we found for Monkey P a distribution that was not significantly unimodal for the perturbation and steady state epochs (Figure 4C). For Monkey M we found a significant unimodal distribution with a major axis near 180° for both epochs (Figure 4H). However, these distributions were qualitatively more dispersed than the distributions from the difference between perturbation and steady-state tuning as indicated by the Rayleigh Statistic (perturbation vs steady-state: R=0.57,0.44; Contra. vs Ipsi. R=0.23, 0.2). These results suggest that directional tuning is much more similar across time epochs than across contralateral and ipsilateral arms.

Although neural activity in M1 was observed for both ipsilateral and contralateral motor tasks, most neurons displayed greater activity for the contralateral limb. From the plane fits we calculated the firing rate for each neuron in its preferred load direction during contralateral and ipsilateral loads. Figure 5A and E compare the firing rate of each neuron during the perturbation epoch from Monkey P and M, respectively. More than 70% of the neurons had a larger firing rate for contralateral loads (median of all neurons Monkey P: 39 sp/s, Monkey M: 24sp/s) than ipsilateral loads (Monkey P: 16 sp/s, Monkey M: 10sp/s). The distributions generated from the difference between each neuron’s contralateral and ipsilateral firing rate was significantly shifted to the right for both monkeys (Figure 5B and F, Wilcoxon signed-rank test Monkey P: z=5.1 p<0.001, Monkey M: z=5 p<0.001) indicating contralateral responses were larger than ipsilateral responses. Similarly, during the steady-state epoch, we found the contralateral loads evoked larger firing rates (Figure 5C and D note scales are smaller than for 5A and B; Monkey P: 24.1 sp/s, Monkey M: 13.3sp/s) than ipsilateral loads (Monkey P: 9.7 sp/s, Monkey M: 9.2sp/s). The distribution of the difference between contralateral and ipsilateral magnitudes were significantly shifted to the right for both monkeys (Figure 5G and H, Wilcoxon signed-rank test Monkey P: z=5.9 p<0.001, Monkey M: z=4.6 p<0.001). However, note that approximately a quarter of neurons displayed larger responses for the ipsilateral arm across epochs and monkeys.

**Figure 5:**
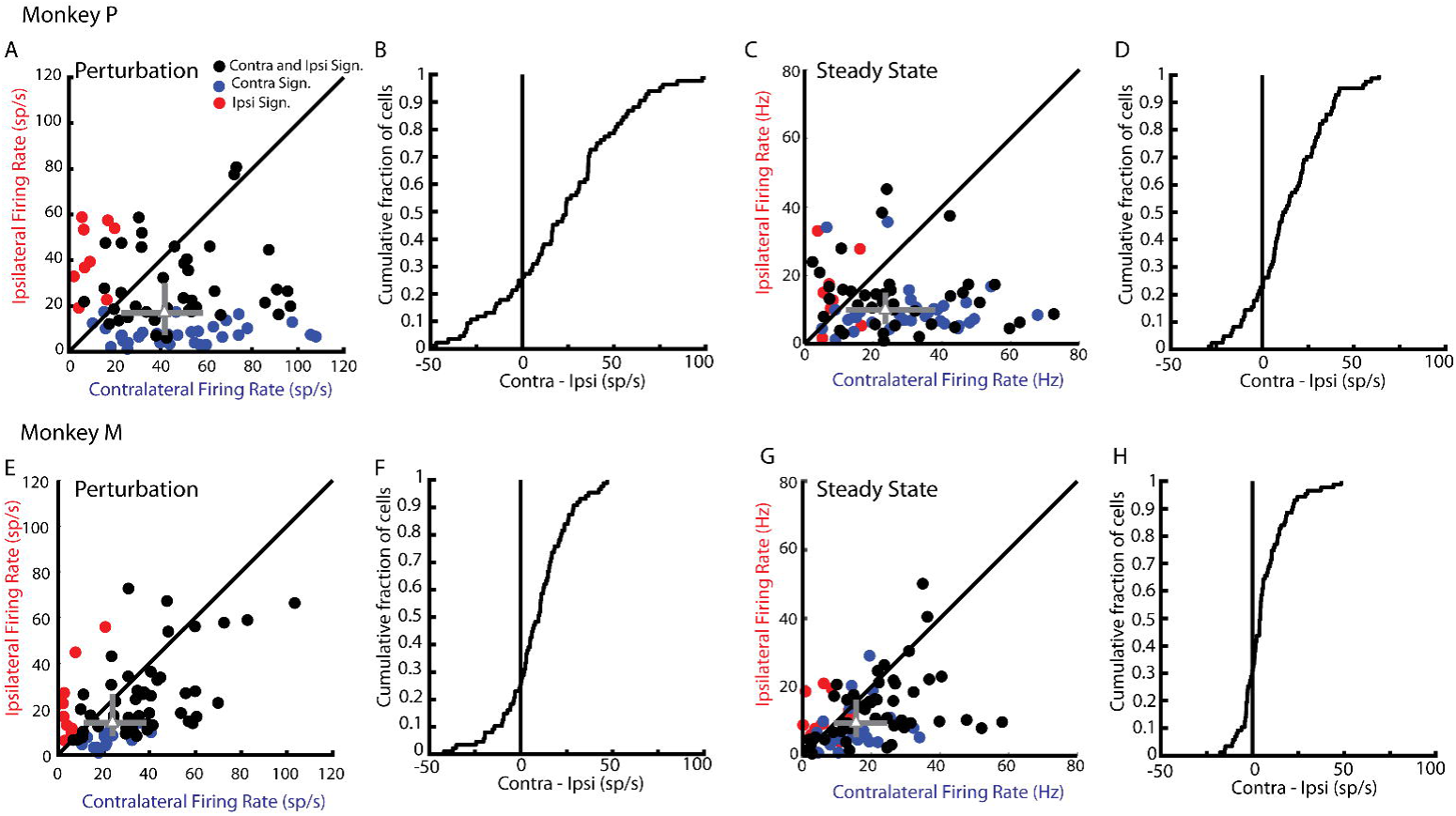
Magnitude comparison between contralateral and ipsilateral neural responses. A) For Monkey P, comparison between the contralateral and ipsilateral firing rates during the perturbation epoch for each neuron as determined by their planar fit. Black circles are neurons that had a significant planar fit for contralateral and ipsilateral loads. Blue and red circles are neurons that had a significant planar fit for contralateral or ipsilateral loads only, respectively. Grey triangle represents the median and the 25^th^ and 75^th^ percentiles. B) The cumulative distribution generated from the difference between contralateral and ipsilateral firing rates. C-D) Same as A-B for steady state. E-H) Same as A-D for Monkey M.

We also examined the onset time of the load response following the perturbation. Figure 6A and D display the averaged change in firing rate across the population of neurons for Monkey P and M for contralateral and ipsilateral loads. For Monkey P, the onset when the firing rate significantly deviated from baseline for contralateral and ipsilateral loads occurred at 28ms and 39ms, respectively. For Monkey M we found onset times that tended to be later than Monkey P with contralateral and ipsilateral loads resulting in changes that started, at 51 and 57ms, respectively, for the population of recorded neurons. Figure 6B and E compare neurons with identified onset times for both contralateral and ipsilateral loads. From this population more than 70% of neurons had an onset time earlier for contralateral (median onset Monkey P: 68ms, Monkey M: 79ms) than ipsilateral loads (Monkey P: 81ms, Monkey M: 117ms). The distribution of the difference between contralateral and ipsilateral onsets were significantly shifted to the left for both monkeys (Wilcoxon signed-rank test Monkey P: z=2.7 p<0.01, Monkey M: z=3.4 p<0.001), indicating contralateral responses tended to be earlier than ipsilateral responses. However, 30% of neurons had response times that were earlier for the ipsilateral arm.

**Figure 6:**
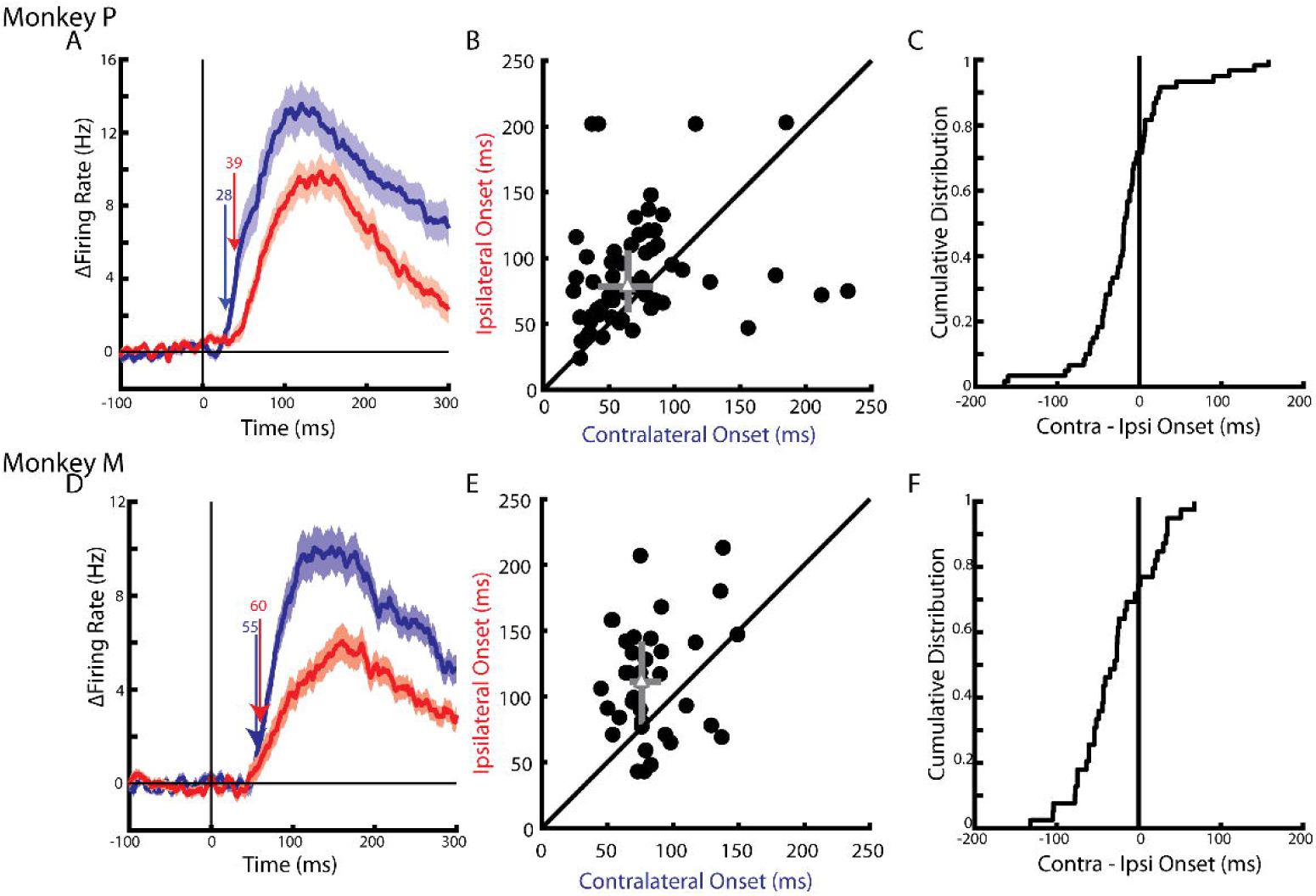
Timing of neural responses to contralateral and ipsilateral loads. A) The average change in firing rate across the population during the contralateral and ipsilateral loads. Arrows mark the onset when a significant change in baseline was detected. B) Comparison of the onsets when contralateral and ipsilateral loads were applied. Grey triangle marks the median and 25^th^ and 75^th^ percentiles. C) The cumulative distribution generated from the difference between contralateral and ipsilateral onset times. D-F) Same as A-C for Monkey M.

We also ensured our main findings were observed using conventional single electrodes by sampling 34 neurons in Monkey P from the opposite hemisphere of the array. We found 18 neurons were load sensitive and their population response started at 35ms for the contralateral loads and 47ms for the ipsilateral loads (data not shown). We also found the magnitude of the contralateral response was larger than the ipsilateral response for 70% of neurons in the perturbation epoch (Median contralateral 19.6 sp/s, ipsilateral 9.2 sp/s). However, during the posture epoch we found ~50% of neurons with a larger contralateral response (Contralateral 6.4 sp/s, Ipsilateral 4.1 sp/s).

### Control Analysis

One possibility for the observed neural activity when ipsilateral loads were applied is that contralateral limb motion that was transferred through the body. We addressed this by fitting the average neural activity when contralateral loads were applied to the average contralateral hand velocity in the perturbation epoch (data not shown). The resulting mappings (one for each neuron) accounted for a large proportion of neural variance. For Monkey P/M, the median variance accounted for (VAF) was 82/66%, and 84/68% of neurons had a VAF greater than 50%. When we used these mappings to predict the neural activity when ipsilateral loads were applied from the contralateral hand motion, we found a substantial decrease in the VAF. For Monkey P/M we found the median VAF was −0.02/0.02%. Further, 55/44% of neurons had a VAF that was less than zero. For Monkey P/M, only 1/2 neurons had a VAF that was greater than 50%.

The poor fits by these mappings may reflect that a neuron’s mapping to hand motion may change depending on the context. We fit the evoked neural activity when ipsilateral loads were applied to the corresponding contralateral hand motion. For Monkey P/M we found these fits had a median VAF of 57/56%, and 63/56% of neurons had a VAF greater than 50%. However, when we examined the mapping weights for the hand motion, we found for Monkey P/M the median weights for the x and y component of the velocities were 6.7/7.4 and 14/27 times larger than the weights found by fitting the contralateral activity with the contralateral hand motion (paragraph above), implying the neurons were much more sensitive to contralateral limb motion when the limb was not directly engaged in a motor task. However, previous studies highlight that sensitivity of neural responses in M1 to limb motion are reduced when the animal is not engaged in a motor action (Omrani et al., 2014, 2016).

### Population Analysis

Given the substantial change in tuning between the contralateral and ipsilateral activity, we questioned whether motor cortex separates contralateral and ipsilateral activity into orthogonal subspaces. If contralateral and ipsilateral activity reside in separate subspaces, then the correlation between pairs of neurons should change between the two contexts. We focus on the perturbation epoch exclusively given the low firing rate of ipsilateral activity in the steady-state for both monkeys. Figure 7A and H show the correlation matrices generated from the pairwise correlation coefficients between all recorded neurons for Monkey P and M. The columns and rows have been ordered to reveal structure in the contralateral matrix (left). However, using the same ordering of columns and rows on the ipsilateral correlation matrices revealed far less structure (left). Likewise, when we permuted the columns and rows of the ipsilateral matrices to reveal structure (Figure 7B and I) we found the same column and row permutations revealed less structure for the contralateral matrices. Figure 7C and J compares the pairwise correlation coefficients between neurons during the contralateral and ipsilateral loads. The population shows a dispersed pattern indicative that the correlation between neurons change substantially between contralateral and ipsilateral loads. Figure 7D and K show the distribution of the absolute change in pairwise correlation coefficient between contralateral and ipsilateral loads. For monkey P and M, we found the median change in correlation coefficient to be 0.32 and 0.20. These median changes were significantly larger than the median generated by a null distribution comparing how pairwise correlations change across randomly selected trials within a context (bootstrap p<0.001 see Methods).

**Figure 7:**
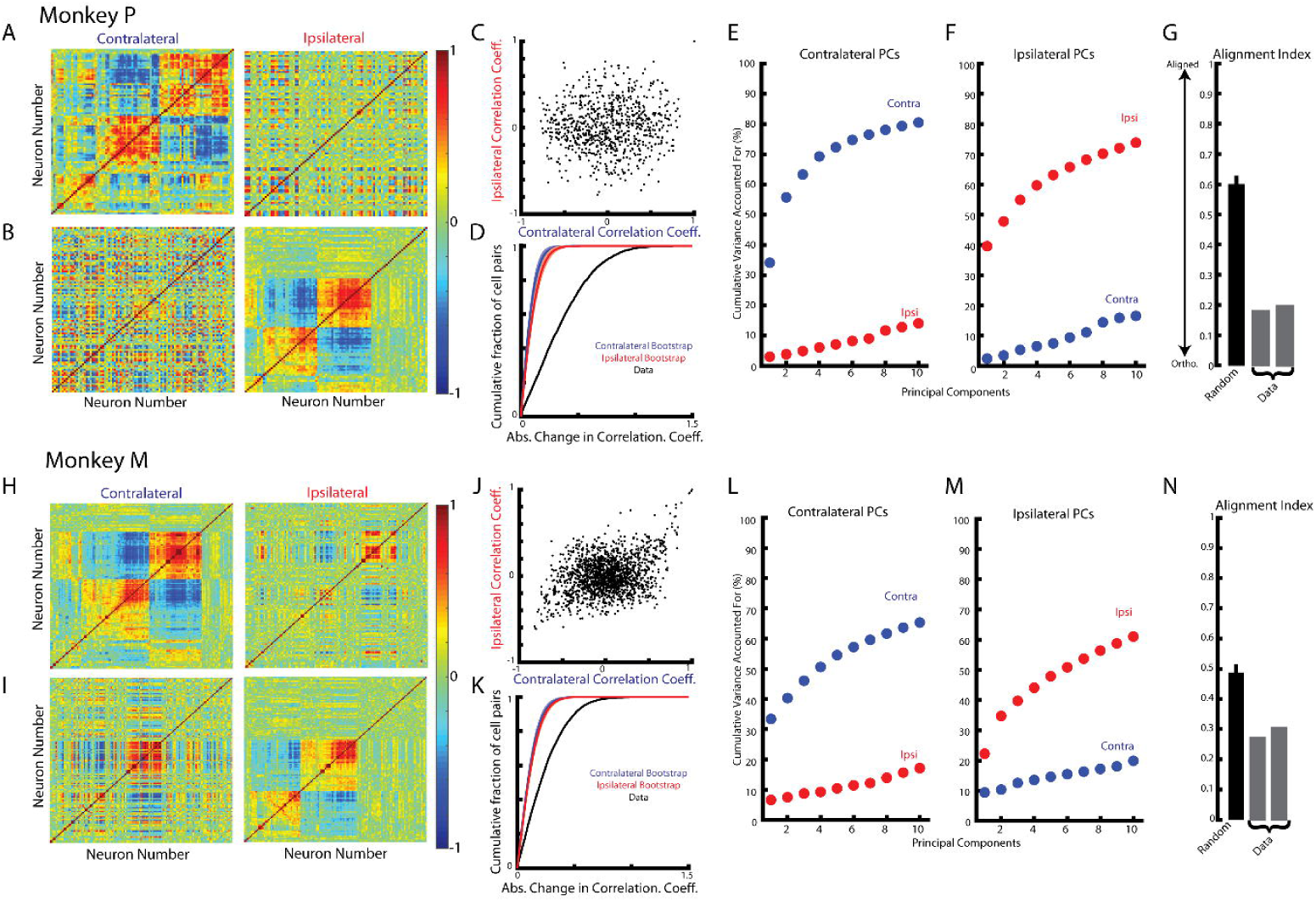
Reorganization of neural population response between contralateral and ipsilateral loads. A) Heatmaps generated from the correlation matrices of the contralateral (left) and ipsilateral activity for Monkey P. The columns and rows of the contralateral matrix were permuted to reveal its structure and the same permutation was applied to the ipsilateral matrix. B) Same as B except the permutations were chosen to reveal structure in the ipsilateral matrix. C) Comparison of the pairwise correlation coefficients between neurons during the contralateral and ipsilateral loads. D) The cumulative sum of the absolute change in the pairwise correlation coefficient between the contralateral and ipsilateral loads (black). A contralateral bootstrap (blue) was generated by separating trials into 2 distinct groups and calculating the absolute change in pairwise correlation coefficient. This was repeated 1000x. Shaded region shows 3 standard deviations from the mean. This was repeated for the ipsilateral data (red). E) The cumulative variance explained of the contralateral and ipsilateral activity from the 10 largest contralateral principle components. F) The cumulative variance explained of the contralateral and ipsilateral activity from the 10 largest ipsilateral principle components. G) The alignment index generated by randomly sampling from the data covariance matrix (black, mean + standard deviation) and the two indices generated from the contralateral and ipsilateral principle components (grey). H-N) Same as A-G for Monkey M.

The substantial change in correlation structure suggests contralateral and ipsilateral activity reside in separate, orthogonal subspaces. Contralateral and ipsilateral subspaces we found using principal component analysis (PCA) which finds linear weightings of neuron responses which captures the largest amount of variance. Figure 7E and L shows the variance captured by the top-ten principle components generated from the contralateral activity. For Monkey P/M, these components captured 84/65% of the contralateral variance, respectively, while accounting for <20% of the ipsilateral variance. Likewise, the top-ten principle components generated from the ipsilateral activity captured 77/61% of the ipsilateral variance but captured ~20% of the contralateral variance (Figure 7F and M). We computed the alignment index to quantify how orthogonal the top contralateral and ipsilateral principle components were (Elsayed et al., 2016). The alignment index can range from 0, indicating perfect orthogonality, to 1 indicating perfect alignment. For Monkey P/M, we found the average alignment index to be 0.19/0.29, respectively (Figure 7G and N). These values were significantly smaller than alignment indices generated by randomly sampling from the data covariance matrix (mean: Monkey P 0.60; Monkey M 0.49; bootstrap p<0.001) indicating contralateral and ipsilateral activity are more orthogonal than expected by random chance.

We observed less variance captured by the top-ten principal components than previous studies (Elsayed et al., 2016; Miri et al., 2017). This may reflect the kernel we used to estimate firing rates as it is not as smooth as kernels used in the previous studies. For comparison with the literature, we re-analyzed our data after convolving with a gaussian kernel (standard deviation 20ms) and found that principal components using the gaussian kernel did explain more variance. We found the top-ten contralateral principle components captured 92/80% of the contralateral variance for Monkey P/M respectively, while capturing <20% of the ipsilateral activity. Similarly, the top-ten ipsilateral principle components captured 90/77% of the ipsilateral variance for Monkey P/M, while capturing <25% of the contralateral variance. The average alignment indices for Monkey P/M were 0.16/0.27 and were significantly smaller than indices generated by randomly sampling from the data covariance (mean: Monkey P 0.67; Monkey M 0.55; bootstrap p<0.001).

One possibility for the separation between contralateral and ipsilateral activity is that PCA is identifying two separate groups of neurons. This seems unlikely given that we observed a substantial proportion of neurons that were active during both contexts. Nonetheless, we addressed this issue following a similar procedure in Perich et al., (2018). We summed the absolute value of the weights from the top ten contralateral (*w_contra_*) and ipsilateral (*w_ipsi_*) principal components. We then calculated the contralateral and ipsilateral difference divided by their sum (*w_contra_* − *w_ipsi_*)/*w_contra_* + *w_ipsi_*). The histogram generated from all neurons did not appear bimodal (data not shown), as would be expected if PCA was identifying separate groups of neurons. Instead we observed a unimodal distribution that was not significantly different from a normal distribution (Kolmogorov-Smirnov test: Monkey P: D(92)=0.06 p=0.9, Monkey M: D(130)=0.08 p=0.3). These data indicate PCA was not simply isolating separate neural populations for contralateral and ipsilateral activity.

Although PCA identified subspaces for contralateral and ipsilateral activity that were close to orthogonal, they were still partially aligned. To identify an orthogonal subspace that captures the contralateral and ipsilateral activity we used a joint optimization procedure from Elsayed et al., (2016). For Monkey P/ M, we found three orthogonal contralateral dimensions that captured 65% /48% of the contralateral variance, which was comparable to the variance captured by the top-three contralateral principle components, 66%/50%, respectively. Projecting the contralateral activity onto these three orthogonal contralateral dimensions reveal time series with substantial time-varying dynamics (Figure 8A and C, left column). By comparison, projecting the ipsilateral activity revealed little change in activity from baseline (right column). We compared the relative difference between the contralateral and ipsilateral time series in the contralateral dimensions (Equation 4 Methods). We found a significant difference across all three dimensions (bootstrap p<0.001) with average difference of 99% /104% for Monkey P/M.

**Figure 8:**
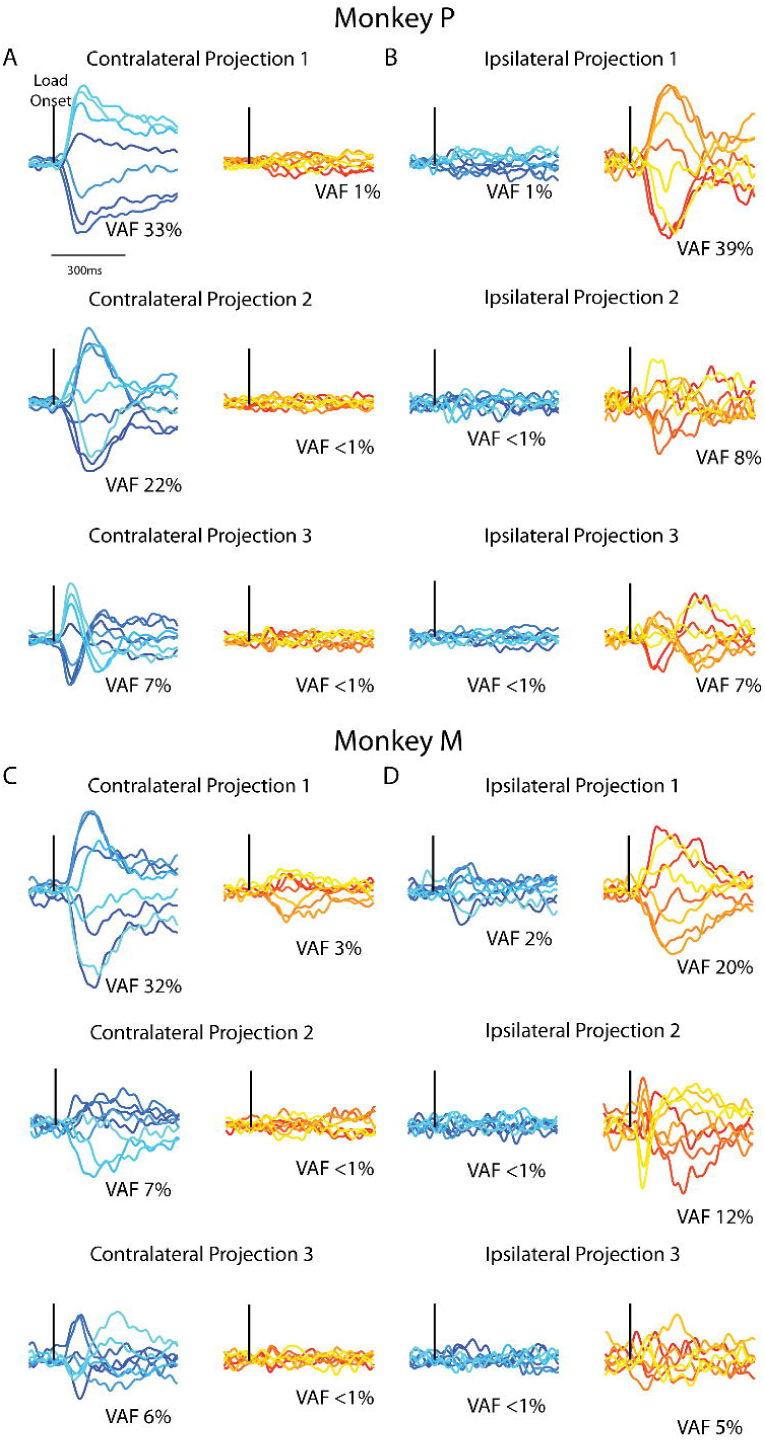
Time series for the orthogonal contralateral and ipsilateral dimensions. A) For Monkey P, the time series generated by projecting the contralateral (left) and ipsilateral (right) activity onto the 3 orthogonal contralateral dimensions. Black line indicates when the load was applied. B) Same as A for the 3 ipsilateral dimensions. C-D) Same A-B for Monkey M.

The three orthogonal ipsilateral dimensions captured 57% /40% of the ipsilateral variance for Monkey P/M and was comparable to the top three ipsilateral principle components, 58% /43%. Examining the ipsilateral activity in the three orthogonal ipsilateral dimensions revealed substantial time-varying dynamics (Figure 8B and C left columns) while contralateral activity in these dimensions changed little from baseline (right columns). The relative difference in amplitudes between the ipsilateral and contralateral dimensions was also significant (p<0.001) with an average difference of 105% /116% for Monkey P/M. These time-series reveal that linearly summing neural responses can isolate the contralateral neural activity from the ipsilateral activity, and vice versa.

## DISCUSSION

Several studies have examined neural responses to the ipsilateral and contralateral limbs during goal-directed reaching (Donchin et al., 1998, 2001, 2002; Kermadi et al., 1998; Cardoso de Oliveira et al., 2001; Gribova et al., 2002; Steinberg et al., 2002; Cisek et al., 2003). We examined how load-related activity for the contralateral and ipsilateral limb are represented using a postural perturbation task. We demonstrate that each neuron’s preferred load has little relationship between the contralateral and ipsilateral limb. We also found the contralateral responses were twice as large and started ~10ms earlier than the ipsilateral responses. Furthermore, we demonstrate that the ipsilateral and contralateral activities exist in separate subspaces, allowing the contralateral and ipsilateral responses to be separated by a weighted sum of each neuron’s response.

An early study suggested M1 was largely insensitive to ipsilateral movement (Tanji et al., 1988). In contrast, we found ~55% of load-sensitive neurons in M1 were sensitive to ipsilateral and contralateral movement in agreement with follow-up studies (Kermadi et al., 1998; Donchin et al., 2002; Cisek et al., 2003). We also found the earliest response to a contralateral load occurred ~10ms sooner than the response for an ipsilateral load, suggesting the corpus callosum is the likely source of ipsilateral information, at least for the fastest ipsilateral responses (Aboitiz et al., 1992; Ringo et al., 1994; Fendrich et al., 2004; Caminiti et al., 2013). This 10ms delay between contralateral and ipsilateral limb matches the additional delay in task-dependent feedback responses between the two limbs during bimanual tasks as compared to unimanual tasks (Marsden et al., 1981; Mutha and Sainburg, 2009; Dimitriou et al., 2012; Omrani et al., 2013).

Although we did observe contralateral limb movement when the ipsilateral limb was perturbed, it is unlikely to be the main cause of the corresponding neural activity. We have previously shown that M1 responses are less sensitive to perturbation-caused limb motion if the limb is not engaged in a task (Omrani et al., 2014, 2016). This contradicts our finding that the weights between contralateral limb motion and evoked neural responses increase (i.e. more sensitive) for ipsilateral perturbations when the contralateral limb was not directly engaged in the task. Secondly, we found negligible changes in muscle activity from baseline during the ipsilateral perturbations. Lastly, during steady-state when contralateral limb movement would have ceased, we still observed significant ipsilateral neural activity.

As in our previous studies (Cabel et al., 2001; Herter et al., 2009), we found the preferred load directions of M1 neurons were not uniformly distributed for the contralateral limb. Instead we found a higher proportion of neurons maximally active for combined elbow flexors and shoulder extensors (whole-limb flexors) and combined elbow extensors and shoulder flexors (whole-limb extensors). Previous work highlights how this pattern of activity parallels the distribution of preferred load directions of proximal limb muscles (Kurtzer et al., 2006). Neural network models that are trained to control a two-joint limb display a similar bimodal pattern of activity across the network, but only when bi-articular muscles representing the biceps and triceps are included in the limb model (Lillicrap and Scott, 2013). Notably, the bias in the distribution is opposite to the anatomical action of the bi-articular muscles (combined flexion for biceps and combined extension for triceps).

Interestingly, the population tuning distribution of M1 activity related to the ipsilateral limb had a similar bias towards whole-limb flexors and whole-limb extensors. Thus, load-related activity related to the ipsilateral limb is not simply an abstract representation of loads. Rather ipsilateral activity reflects the mechanical and anatomical properties of the limb musculature even though the descending projection from this hemisphere predominantly target the contralateral musculature.

Cisek et al., (2003) demonstrated that the correlation in the preferred direction of M1 neurons during reaching between contralateral and ipsilateral limbs can vary anatomically. Areas more rostral exhibited a high correlation while more caudal areas exhibit low correlation between the contralateral and ipsilateral preferred direction. In our study, we found Monkey P exhibited no correlation between the preferred load for the contralateral and ipsilateral limb, whereas for Monkey M we found a small correlation. This may indicate that the array for Monkey M was placed more rostral than for Monkey P.

The fact that neurons can represent different loads for the contralateral and ipsilateral limb appears to be at odds with a recent study which used fMRI imaging to examine areas of M1 during a finger pressing task (Diedrichsen et al., 2013). They found that voxels in M1 preferred similar finger representations in both hands. It may be that the organization of M1 related to the proximal and distal limbs are distinct. Alternatively, their analysis may have been primarily dominated by rostral M1 as voxel-based analysis aggregates activity associated with a large number of neurons and may not be able to observe this level of cortical organization.

We also found neural activity related to each limb could be isolated into orthogonal subspaces. A property of orthogonality is that by summing a linear weighting of each neuron’s responses, we can isolate the contralateral activity from the ipsilateral activity. This may provide a mechanism for how ipsilateral activity does not generate motor output. The synaptic weights of M1 neurons onto the spinal cord circuits could be assigned such that the net effect of ipsilateral activity cancel out. This has been proposed for how preparatory activity in motor cortex does not cause motor output (Kaufman et al., 2014; Elsayed et al., 2016).

Likewise, brain regions that utilize the ipsilateral activity can weight each neuron’s response to extract the ipsilateral activity without interference from the contralateral activity. Li et al., (2016) highlights how activity in the ipsilateral M1 improves the robustness of activity in the contralateral M1. The ipsilateral subspace may also allow for flexible bimanual coupling when both limbs are utilized for the same task goal (Marsden et al., 1981; Diedrichsen, 2007; Dimitriou et al., 2012; Omrani et al., 2013). This ability to segregate different pieces of information from neural activity by summing the weighted activity of neurons has been observed across many different brain regions (Pouget and Sejnowski, 1997; Stopfer et al., 2003; Machens et al., 2010; Mante et al., 2013; Raposo et al., 2014; Kobak et al., 2016; Gallego et al., 2017; Michaels et al., 2018; Remington et al., 2018).

Orthogonality may also allow different spinal circuits to be engaged depending on the behaviour. Miri et al., (2017) found motor cortex was involved in modulating reaching and locomotive motor output, however with different latencies suggesting motor cortex was engaging separate spinal cord circuits. Consistent with this hypothesis, activity in motor cortex during reaching was orthogonal to the activity for locomotion. A similar mechanism may exist for load-related activity during posture and reaching as we have previously found no relationship between a neuron’s gain between these two contexts that may reflect an orthogonal subspace (Kurtzer et al., 2005).

Also, our finding of orthogonality between contexts is not trivial. Recent studies show neural activity remains within a subspace after learning a curl-force field (Perich et al., 2018), visuomotor rotation (Perich et al., 2018; Vyas et al., 2018), or BMI mappings (Golub et al., 2018) and across reaching tasks with different load conditions (Gribble and Scott, 2002; Gallego et al., 2018) and initiation cues (Lara et al., 2018).

Recently, Ames and Churchland, (2019) compared M1 responses to contralateral and ipsilateral cycling movements. Similar to our findings, they observed contralateral activity that spanned a subspace that was orthogonal to the subspace ipsilateral activity spanned. Furthermore, they found ipsilateral M1 activity could reliably decode ipsilateral muscle activity suggesting ipsilateral activity was reflecting low-level features of the motor output.

The fact that neural activity may be maintained in a subspace that does not influence descending output at the spinal level has implications for a long-standing debate regarding how M1 is involved in descending control. Many studies have found activity in M1 correlates with muscle activity (Evarts, 1968; Humphrey, 1972; Murphy et al., 1985; Bennett and Lemon, 1996; Scott, 1997; Sergio and Kalaska, 1998; Kakei et al., 1999; Cherian et al., 2013; Oby et al., 2013), suggesting that M1 activity reflected low-level features of the motor output such as the spatio-temporal features of muscle activity. However other studies have correlated M1 activity with whole limb movements (Georgopoulos et al., 1982), trajectories (Schwartz, 1992, 1993) and other high level parameters of motion such as target direction and speed (Johnson et al., 1999). Even when high and low-level features of motor action are dissociated, some activity still appears to be related to high level features of the movement (Thach, 1978; Sergio and Kalaska, 1998; Kakei et al., 1999; Russo et al., 2018). Our data suggests that both sets of information could be present in M1 simultaneously but only low-level motor commands to muscles influences spinal circuits. By retaining different types or sources of information in different subspaces, neural activity can both control motor output and process other types of information, simultaneously.

## REFERENCES

Aboitiz F, Scheibel AB, Fisher RS, Zaidel E (1992) Fiber composition of the human corpus callosum. Brain Res 598:143–153.

Ames KC, Churchland MM (2019) Motor cortex signals corresponding to the two arms are shared across hemispheres, mixed among neurons, yet partitioned within the population response. bioRxiv Available at: http://biorxiv.org/lookup/doi/10.1101/552257 [Accessed March 18, 2019].

Batschelet E (1981) Circular Statistics in Biology. New York: Academic Press.

Bennett KM, Lemon RN (1996) Corticomotoneuronal contribution to the fractionation of muscle activity during precision grip in the monkey. J Neurophysiol 75:1826–1842.

Boumal N, Mishra B, Absil PA, Sepulchre R (2014) Manopt, a Matlab Toolbox for Optimization on Manifolds. J Mach Learn Res 15:1455–1459.

Brösamle C, Schwab ME (1997) Cells of origin, course, and termination patterns of the ventral, uncrossed component of the mature rat corticospinal tract. J Comp Neurol 386:293–303.

Cabel DW, Cisek P, Scott SH (2001) Neural Activity in Primary Motor Cortex Related to Mechanical Loads Applied to the Shoulder and Elbow During a Postural Task. J Neurophysiol 86:2102–2108.

Caminiti R, Carducci F, Piervincenzi C, Battaglia-Mayer A, Confalone G, Visco-Comandini F, Pantano P, Innocenti GM (2013) Diameter, Length, Speed, and Conduction Delay of Callosal Axons in Macaque Monkeys and Humans: Comparing Data from Histology and Magnetic Resonance Imaging Diffusion Tractography. J Neurosci 33:14501–14511.

Cardoso de Oliveira S, Gribova A, Donchin O, Bergman H, Vaadia E (2001) Neural interactions between motor cortical hemispheres during bimanual and unimanual arm movements. Eur J Neurosci 14:1881–1896.

Cherian A, Fernandes HL, Miller LE (2013) Primary motor cortical discharge during force field adaptation reflects muscle-like dynamics. J Neurophysiol 110:768–783.

Churchland MM, Cunningham JP, Kaufman MT, Foster JD, Nuyujukian P, Ryu SI, Shenoy KV (2012) Neural population dynamics during reaching. Nature 487:51–56.

Cisek P, Crammond DJ, Kalaska JF (2003) Neural activity in primary motor and dorsal premotor cortex in reaching tasks with the contralateral versus ipsilateral arm. J Neurophysiol 89:922–942.

Cramer SC, Finklestein SP, Schaechter JD, Bush G, Rosen BR (1999) Activation of Distinct Motor Cortex Regions During Ipsilateral and Contralateral Finger Movements. J Neurophysiol 81:383–387.

Cunningham JP, Ghahramani Z (2015) Linear Dimensionality Reduction: Survey, Insights, and Generalizations. J Mach Learn Res 16:2859–2900.

Diedrichsen J (2007) Optimal Task-Dependent Changes of Bimanual Feedback Control and Adaptation. Curr Biol 17:1675–1679.

Diedrichsen J, Wiestler T, Krakauer JW (2013) Two Distinct Ipsilateral Cortical Representations for Individuated Finger Movements. Cereb Cortex 23:1362–1377.

Dimitriou M, Franklin DW, Wolpert DM (2012) Task-dependent coordination of rapid bimanual motor responses. J Neurophysiol 107:890–901.

Donchin O, Gribova A, Steinberg O, Bergman H, Oliveira CS de, Vaadia E (2001) Local field potentials related to bimanual movements in the primary and supplementary motor cortices. Exp Brain Res 140:46–55.

Donchin O, Gribova A, Steinberg O, Bergman H, Vaadia E (1998) Primary motor cortex is involved in bimanual coordination. Nature 395:274–278.

Donchin O, Gribova A, Steinberg O, Mitz AR, Bergman H, Vaadia E (2002) Single-Unit Activity Related to Bimanual Arm Movements in the Primary and Supplementary Motor Cortices. J Neurophysiol 88:3498–3517.

Druckmann S, Chklovskii DB (2012) Neuronal Circuits Underlying Persistent Representations Despite Time Varying Activity. Curr Biol 22:2095–2103.

Dum RP, Strick PL (1996) Spinal Cord Terminations of the Medial Wall Motor Areas in Macaque Monkeys. J Neurosci 16:6513–6525.

Elsayed GF, Lara AH, Kaufman MT, Churchland MM, Cunningham JP (2016) Reorganization between preparatory and movement population responses in motor cortex. Nat Commun 7 Available at: http://www.nature.com/articles/ncomms13239 [Accessed November 26, 2018].

Evarts EV (1968) Relation of pyramidal tract activity to force exerted during voluntary movement. J Neurophysiol 31:14–27.

Fendrich R, Hutsler J, Gazzaniga M (2004) Visual and tactile interhemispheric transfer compared with the method of Poffenberger. Exp Brain Res 158 Available at: http://link.springer.com/10.1007/s00221-004-1873-6 [Accessed February 25, 2019].

Gallego JA, Perich MG, Miller LE, Solla SA (2017) Neural Manifolds for the Control of Movement. Neuron 94:978–984.

Gallego JA, Perich MG, Naufel SN, Ethier C, Solla SA, Miller LE (2018) Cortical population activity within a preserved neural manifold underlies multiple motor behaviors. Nat Commun 9 Available at: http://www.nature.com/articles/s41467-018-06560-z [Accessed November 16, 2018].

Gallivan JP, McLean DA, Smith FW, Culham JC (2011) Decoding Effector-Dependent and Effector-Independent Movement Intentions from Human Parieto-Frontal Brain Activity. J Neurosci 31:17149–17168.

Georgopoulos AP, Kalaska JF, Caminiti R, Massey JT (1982) On the relations between the direction of two-dimensional arm movements and cell discharge in primate motor cortex. J Neurosci 2:1527–1537.

Golub MD, Sadtler PT, Oby ER, Quick KM, Ryu SI, Tyler-Kabara EC, Batista AP, Chase SM, Yu BM (2018) Learning by neural reassociation. Nat Neurosci 21:607–616.

Gribble PL, Scott SH (2002) Overlap of internal models in motor cortex for mechanical loads during reaching. Nature 417:938–941.

Gribova A, Donchin O, Bergman H, Vaadia E, Oliveira SC de (2002) Timing of bimanual movements in human and non-human primates in relation to neuronal activity in primary motor cortex and supplementary motor area. Exp Brain Res 146:322–335.

Heming EA, Lillicrap TP, Omrani M, Herter TM, Pruszynski JA, Scott SH (2016) Primary motor cortex neurons classified in a postural task predict muscle activation patterns in a reaching task. J Neurophysiol 115:2021–2032.

Herter TM, Korbel T, Scott SH (2009) Comparison of Neural Responses in Primary Motor Cortex to Transient and Continuous Loads During Posture. J Neurophysiol 101:150–163.

Humphrey DR (1972) Relating motor cortex spike trains to measures of motor performance. Brain Res 40:7–18.

Johnson MTV, Coltz JD, Ebner TJ (1999) Encoding of target direction and speed during visual instruction and arm tracking in dorsal premotor and primary motor cortical neurons. Eur J Neurosci 11:4433–4445.

Kakei S, Hoffman DS, Strick PL (1999) Muscle and Movement Representations in the Primary Motor Cortex. Science 285:2136–2139.

Kaufman MT, Churchland MM, Ryu SI, Shenoy KV (2014) Cortical activity in the null space: permitting preparation without movement. Nat Neurosci 17:440–448.

Kermadi I, Calciati T, Rouiller EM (1998) Neuronal activity in the primate supplementary motor area and the primary motor cortex in relation to spatio-temporal bimanual coordination. Somatosens Mot Res 15:287–308.

Kobak D, Brendel W, Constantinidis C, Feierstein CE, Kepecs A, Mainen ZF, Qi X-L, Romo R, Uchida N, Machens CK (2016) Demixed principal component analysis of neural population data. eLife 5 Available at: https://elifesciences.org/articles/10989 [Accessed February 21, 2019].

Kobayashi M, Hutchinson S, Schlaug G, Pascual-Leone A (2003) Ipsilateral motor cortex activation on functional magnetic resonance imaging during unilateral hand movements is related to interhemispheric interactions. NeuroImage 20:2259–2270.

Kurtzer I, Herter TM, Scott SH (2005) Random change in cortical load representation suggests distinct control of posture and movement. Nat Neurosci 8:498–504.

Kurtzer I, Herter TM, Scott SH (2006) Nonuniform Distribution of Reach-Related and Torque-Related Activity in Upper Arm Muscles and Neurons of Primary Motor Cortex. J Neurophysiol 96:3220–3230.

Lacroix S, Havton LA, McKay H, Yang H, Brant A, Roberts J, Tuszynski MH (2004) Bilateral corticospinal projections arise from each motor cortex in the macaque monkey: A quantitative study. J Comp Neurol 473:147–161.

Lara AH, Elsayed GF, Zimnik AJ, Cunningham JP, Churchland MM (2018) Conservation of preparatory neural events in monkey motor cortex regardless of how movement is initiated. eLife 7:e31826.

Li N, Daie K, Svoboda K, Druckmann S (2016) Robust neuronal dynamics in premotor cortex during motor planning. Nature 532:459–464.

Lillicrap TP, Scott SH (2013) Preference Distributions of Primary Motor Cortex Neurons Reflect Control Solutions Optimized for Limb Biomechanics. Neuron 77:168–179.

Machens CK, Romo R, Brody CD (2010) Functional, But Not Anatomical, Separation of “What” and “When” in Prefrontal Cortex. J Neurosci 30:350–360.

Mante V, Sussillo D, Shenoy KV, Newsome WT (2013) Context-dependent computation by recurrent dynamics in prefrontal cortex. Nature 503:78–84.

Marsden CD, Merton PA, Morton HB (1981) Human Postural Responses. Brain 104:513–534.

McKiernan BJ, Marcario JK, Karrer JH, Cheney PD (1998) Corticomotoneuronal Postspike Effects in Shoulder, Elbow, Wrist, Digit, and Intrinsic Hand Muscles During a Reach and Prehension Task. J Neurophysiol 80:1961–1980.

Michaels JA, Dann B, Intveld RW, Scherberger H (2018) Neural Dynamics of Variable Grasp-Movement Preparation in the Macaque Frontoparietal Network. J Neurosci 38:5759–5773.

Miri A, Warriner CL, Seely JS, Elsayed GF, Cunningham JP, Churchland MM, Jessell TM (2017) Behaviorally Selective Engagement of Short-Latency Effector Pathways by Motor Cortex. Neuron 95:683–696.e11.

Montgomery LR, Herbert WJ, Buford JA (2013) Recruitment of ipsilateral and contralateral upper limb muscles following stimulation of the cortical motor areas in the monkey. Exp Brain Res 230:153–164.

Murphy JT, Wong YC, Kwan HC (1985) Sequential activation of neurons in primate motor cortex during unrestrained forelimb movement. J Neurophysiol 53:435–445.

Mutha PK, Sainburg RL (2009) Shared Bimanual Tasks Elicit Bimanual Reflexes During Movement. J Neurophysiol 102:3142–3155.

Oby ER, Ethier C, Miller LE (2013) Movement representation in the primary motor cortex and its contribution to generalizable EMG predictions. J Neurophysiol 109:666–678.

Omrani M, Diedrichsen J, Scott SH (2013) Rapid feedback corrections during a bimanual postural task. J Neurophysiol 109:147–161.

Omrani M, Murnaghan CD, Pruszynski JA, Scott SH (2016) Distributed task-specific processing of somatosensory feedback for voluntary motor control. eLife 5:e13141.

Omrani M, Pruszynski JA, Murnaghan CD, Scott SH (2014) Perturbation-evoked responses in primary motor cortex are modulated by behavioral context. J Neurophysiol 112:2985–3000.

Perich MG, Gallego JA, Miller L (2018) A Neural Population Mechanism For Rapid Learning. Neuron 100:964–976.

Pouget A, Sejnowski TJ (1997) Spatial Transformations in the Parietal Cortex Using Basis Functions. J Cogn Neurosci 9:222–237.

Raposo D, Kaufman MT, Churchland AK (2014) A category-free neural population supports evolving demands during decision-making. Nat Neurosci 17:1784–1792.

Remington ED, Narain D, Hosseini EA, Jazayeri M (2018) Flexible Sensorimotor Computations through Rapid Reconfiguration of Cortical Dynamics. Neuron 98:1005–1019.e5.

Ringo JL, Doty RW, Demeter S, Simard PY (1994) Time is of the essence: A conjecture that hemispheric specialization arises from interhemispheric conduction delay. Cereb Cortex 4:331–343.

Rosenzweig ES, Brock JH, Culbertson MD, Lu P, Moseanko R, Edgerton VR, Havton LA, Tuszynski MH (2009) Extensive Spinal Decussation and Bilateral Termination of Cervical Corticospinal Projections in Rhesus Monkeys. J Comp Neurol 513:151–163.

Russo AA, Bittner SR, Perkins SM, Seely JS, London BM, Lara AH, Miri A, Marshall NJ, Kohn A, Jessell TM, Abbott LF, Cunningham JP, Churchland MM (2018) Motor Cortex Embeds Muscle-like Commands in an Untangled Population Response. Neuron 97:953–966.e8.

Schwartz AB (1992) Motor cortical activity during drawing movements: single-unit activity during sinusoid tracing. J Neurophysiol 68:528–541.

Schwartz AB (1993) Motor cortical activity during drawing movements: population representation during sinusoid tracing. J Neurophysiol 70:28–36.

Scott SH (1997) Comparison of Onset Time and Magnitude of Activity for Proximal Arm Muscles and Motor Cortical Cells Before Reaching Movements. J Neurophysiol 77:1016–1022.

Scott SH (1999) Apparatus for measuring and perturbing shoulder and elbow joint positions and torques during reaching. J Neurosci Methods 89:119–127.

Sergio LE, Kalaska JF (1998) Changes in the Temporal Pattern of Primary Motor Cortex Activity in a Directional Isometric Force Versus Limb Movement Task. J Neurophysiol 80:1577–1583.

Smith WS, Fetz EE (2009) Synaptic Linkages Between Corticomotoneuronal Cells Affecting Forelimb Muscles in Behaving Primates. J Neurophysiol 102:1040–1048.

Soteropoulos DS, Edgley SA, Baker SN (2011) Lack of Evidence for Direct Corticospinal Contributions to Control of the Ipsilateral Forelimb in Monkey. J Neurosci 31:11208–11219.

Stavisky SD, Kao JC, Ryu SI, Shenoy KV (2017) Motor Cortical Visuomotor Feedback Activity Is Initially Isolated from Downstream Targets in Output-Null Neural State Space Dimensions. Neuron 95:195–208.e9.

Steinberg O, Donchin O, Gribova A, De Oliveira SC, Bergman H, Vaadia E (2002) Neuronal populations in primary motor cortex encode bimanual arm movements. Eur J Neurosci 15:1371–1380.

Stopfer M, Jayaraman V, Laurent G (2003) Intensity versus Identity Coding in an Olfactory System. Neuron 39:991–1004.

Tanji J, Okano K, Sato KC (1988) Neuronal activity in cortical motor areas related to ipsilateral, contralateral, and bilateral digit movements of the monkey. J Neurophysiol 60:325–343.

Thach WT (1978) Correlation of neural discharge with pattern and force of muscular activity, joint position, and direction of intended next movement in motor cortex and cerebellum. J Neurophysiol 41:654–676.

Thompson KG, Hanes DP, Bichot NP, Schall JD (1996) Perceptual and motor processing stages identified in the activity of macaque frontal eye field neurons during visual search. J Neurophysiol 76:4040–4055.

Vyas S, Even-Chen N, Stavisky SD, Ryu SI, Nuyujukian P, Shenoy KV (2018) Neural Population Dynamics Underlying Motor Learning Transfer. Neuron 97:1177–1186.e3.

